# Leptospirosis in Campinas, Brazil: The interplay between drainage, impermeable areas, and social vulnerability

**DOI:** 10.1101/2024.12.10.627826

**Authors:** Thiago Salomão de Azevedo, Shahista Nisa, Stuart Littlejohn, Renata L. Muylaert

## Abstract

Leptospirosis is an epidemic disease caused by bacteria of the *Leptospira* genus. Its risk is closely associated with inadequate sanitation and flooding, a common public health challenge in large urban centers together with urban environmental modifications, and socio-economic factors. This retrospective observational research investigated the association between the distribution of leptospirosis cases and three contextual factors —drainage, soil impermeability and social vulnerability—in Campinas city, São Paulo, Brazil. We hypothesized that the number of cases will increase in areas that are impermeable and in proximity to drainage systems as well as where social vulnerability is high. We investigated the associations based on 83 autochthonous cases, comparing cases where infection risk was linked to contact with floodwater or mud (n=54) to cases associated with other exposures (n=29). Spatial statistics were used to map disease distribution and investigate the relationship between leptospirosis cases and contextual factors. Our results indicate that the density of leptospirosis increases near drainage systems with risk peaking at 200 m, in areas of greater social vulnerability with increased risk due to floodwater or mud exposure in highly vulnerable regions, and in highly waterproofed urban areas. This study demonstrated that leptospirosis risk remains highly determined by living and working conditions. These findings support targeted strategies to deliver effective prevention, treatment and control interventions in highly populated urban areas of the Global South and similar contexts. Furthermore, combining local contextual environmental information with spatial analysis produces relevant evidence for guiding health public policy and spatial planning and provides precise parameters for future epidemiological models and prevention actions.

**Author summary:** This study explores the links between environmental and contextual factors that influence the risk of leptospirosis transmission to humans in Campinas, São Paulo, Brazil. Leptospirosis is an infectious disease caused by the *Leptospira* bacteria. We investigated how drainage systems, impermeable soil area, and social vulnerability contribute to disease distribution, using spatial statistics to map spatial conditions for 83 cases associated with water and mud contact and other risks. Our findings highlight how environmental and socio-economic factors intersect to influence public health, shedding light on the role of urban planning and drainage infrastructure in the transmission risk of infectious diseases. This interdisciplinary approach underscores the importance of considering social and environmental contexts when developing public health strategies, aligning with broader global efforts to address diseases linked to urbanization and environmental changes. Our study advances the understanding of how spatial data and environmental factors can guide more precise parameters for epidemiological models, offering insights into disease control interventions. We discuss the role of prevention, flood management, and equitable infrastructure in safeguarding public health, emphasizing how the environment shapes health risks. This research provides practical recommendations for helping decision-makers prioritize areas for intervention to reduce the burden of leptospirosis, particularly in vulnerable communities.

## Introduction

Public health in urban areas faces different hazards associated with suboptimal urban planning, social vulnerability and climate-related hazards [1]. A hazard that is likely to increase is flooding, an extreme phenomenon that severely impacts human populations. Flooding increases the risk of infectious and non-communicable diseases, including leptospirosis [2,3], hepatitis [4], mental illness and heart problems [5]. Leptospirosis has been recognized as an infection of intensifying global public health concern and particularly in the Global South [6], yet it remains a neglected tropical disease [7]. Annually, more than one million symptomatic cases, resulting in at least 60,000 deaths, are reported worldwide [8]. The symptoms of leptospirosis in humans are usually non-specific, leading to frequent misdiagnosis or delays in diagnosis. The classic presentation includes fever, chills, severe headache, muscle aches, conjunctival suffusion, nausea, and vomiting [9]. Severe forms of leptospirosis are Weil’s disease, with jaundice and renal insufficiency or interstitial alveolar hemorrhage, congestion, and edema, and pulmonary hemorrhage syndrome, which is a severe lung disease [10,11]. The disease causes a heavy health burden in tropical regions, particularly in areas characterized by poverty, humid climate, and lack of adequate health infrastructures [12].

Leptospirosis is caused by infection from bacteria of the genus *Leptospira*, which can infect land and sea mammals [13], including domestic animals, wildlife, and humans [14]. Human infections occur from direct contact with infected urine and excretions by mammals [15] or indirectly through the contaminated environment [16]. The greatest risk factor for human infection is exposure to contaminated water [17]. Water offers a suitable medium for leptospires, agglutinated with organic matter, to remain infectious for long periods of time [14]. This risk is amplified in urban settings, where high population density and unpredictable flooding can increase exposure and hinder effective protection. Rats are the main hosts in urban settings which, combined with water, creates ideal conditions for leptospirosis transmission as leptospires can survive in water and humid soil [18]. Occupation and poverty can increase leptospirosis risk in areas where there is pervasive bad sanitation, rodent infestation, flooded soils, and lack of access to healthcare. However, the precise magnitude and combined effects of these factors remain unknown for many cities. For instance, exposure may vary with social vulnerability and geographical location, underscoring the need for studies that map cases together with social and environmental exposure factors. Such quantified data can help allocate surveillance and prevention efforts in urban infrastructure.

Recent decades have seen an increase in the incidence of leptospirosis in urban areas prone to inundation [19]. Many cities are designed and built in proximity with water bodies because the resources these environments provide are essential for population livelihood, particularly during city formation [20]. However, several anthropic activities that promote contact with these sites lead to changes in the biological cycles of disease vectors, hosts and reservoirs, as well as in the ways humans are exposed to them [21]. In Brazil, the growing urbanization in large cities contributes to leptospirosis outbreaks because the concentration of human habitat generates residues that can serve as refuge and provide resources for rodents [3]. In addition, the increase in urban sprawling and paved roads and streets cause soil impermeability and prevent or hinder the infiltration of rainwater, resulting in flooding and a greater number of infected people [20,22]. Populations of low socioeconomic status are more affected by leptospirosis due to the synergistic effect of factors such as low-quality sanitation services, environmental and housing characteristics and work activities that often favor contact with the vector [23,24].

In geomorphology, drainage systems, also known as river systems, refer to the structural patterns formed by streams, rivers, and lakes within a drainage basin. Because of gravity and variations in soil permeability, drainage patterns range from dendritic or fractal to paralel and rectangular. These patterns depend highly on the topography and geology of the land, and include artificial alterations. In urban settings, drainage involves either the natural or artificial removal of surface and subsurface water from an area with excess water through ditches, plantings, drain gutters and channeled rivers. In such settings, we expect that the ideal condition for leptospirosis transmission via contaminated water during a flood happens when two conditions are met. First, areas will be closer to drainage systems and second, these areas will have high soil impermeability.

As a thorough understanding of the ecology of leptospirosis in relation with urban drainage and social vulnerability are important for the control of outbreaks, this study utilizes Geographic Information Systems (GIS) tools and spatial analysis to identify the geographical factors associated with leptospirosis occurrence in Campinas, São Paulo, Brazil. We hypothesize that areas with critical flood points have a higher frequency leptospirosis cases, as they meet two key flood-prone risk conditions: proximity to drainage systems and high levels of impermeability to water. Additionally, we expect a negative correlation between leptospirosis incidence and socioeconomic status, with cases concentrated in areas of greater social vulnerability. To test this hypothesis, we focus on cases where infection risk is linked to contact with floodwater or mud, comparing them to cases associated with other exposures. Our primary objective was to quantify spatial patterns of the disease, delineate high-risk areas, and assess environmental factors contributing to its spread. By doing so, we aim to support epidemiological surveillance efforts and enhance the response to health emergencies related to leptospirosis.

## Methods

### Study Area

The municipality of Campinas is located in the interior of São Paulo State with an estimated population of 1,139,047 inhabitants and a population density of 1,433.541 hab / km² [25]. The climate is characterized as Tropical at altitude (Cwa) according to the Koppen / Geiger classification [26]. The annual temperature average is 20.7 °C, with dry and mild winters and a long, rainy summer [27] .

### Leptospirosis Data

The incubation period for leptospirosis ranges from two to 30 days, according to the Brazilian Health Surveillance Manual. The notification form of the Information System for Notifiable Diseases (SINAN) [28] for leptospirosis contains a section in which the risk factors for the disease are primarily associated with the environment and the patients’ exposure conditions [29]. The main risk factors in the Additional Case Data section are listed in Table 1.

**Table 1.** Associated risks reported in the notification form in the Information System for Notifiable Diseases (SINAN) for leptospirosis in Brazil.

| Risk | Description |
| --- | --- |
| Risk related to floodwater or mud | Exposure to floodwater or mud from floods is one of the primary risk factors for infection; |
| Contact with rodents | Direct contact with rodents or their excrement and urine is a common form of leptospirosis transmission and the presence of rodent signs in areas where the patient lives or works is a significant risk factor, as these animals can transmit the bacteria through their urine. |
| Vacant lots | Contact with undeveloped or abandoned land can be a source of infection due to the presence of rodents and other animals. |
| Contact with garbage | Handling waste or working in places with accumulated trash can also be a risk situation. |
| Animal husbandry | The presence of domestic or farm animals (such as pigs, cows, etc.) that can be hosts of leptospirosis is a risk factor. |
| Cleaning water tanks | Exposure to untreated water from water tanks can be a source of contamination. |
| Exposure to sewage | Contact with sewage, which may be contaminated with the leptospirosis bacteria, is also a risk situation. |
| Contact with grain storage | Contact with stored grains that may have been contaminated by rodents or other animals. |
| Exposure to bodies of water | Swimming or coming into contact with rivers, lakes, or other bodies of water that could be contaminated. |
| Contact with crops | Contact with harvested crops or stored grains that may have been contaminated by rodents or other animals. |
| Others | Other risks not defined previously. |

These risk factors are recorded in the notification form and help public health authorities investigate the origin of cases and identify high-risk areas for leptospirosis infection, allowing for more effective preventive measures.

Thus, patients with well-defined symptoms who had contact with any risk factor within this time window were considered target cases in our retrospective observational study. The anonymized database of autochthonous cases of leptospirosis from 2007 to 2010 was made available by the Campinas Municipal Health Department. Anonymized patient variables included only the cases that were laboratory confirmed, through the IGM - ELISA method or by Microscopic Agglutination, by Epidemiological Surveillance. These were georeferenced for the local probable occurrence on each record on the urban cadastral map. The yearly number of cases from 2001 to 2023 were incorporated to reveal disease dynamics across time based on SINAN data [28]. Yearly leptospirosis cases were plotted using R 4.4.1 [30] ang *ggplot* [31].

### Spatial distribution of cases and environmental critical points

The verification of rainfall data and anthropogenic variables in the urban area of Campinas, São Paulo, was carried out based on the application of a sensitivity and spatial analysis. Standardized regression coefficients (SRC) and regression models were applied to assess the impact of these variables on leptospirosis cases. A high absolute value for the coefficient indicates that the input variable has a strong influence on leptospirosis risk. An SRC close to zero suggests that the variable has little influence on the model. To perform this procedure, a spatial database containing the following variables was first created: accumulated precipitation on the day that caused the flood event, the São Paulo social vulnerability index (IPVS), and the impermeable area index. Then, we calculated the distance of cases to the closest drainage system, to quantify the spatial relationship between disease occurrence and drainage. We defined drainage as any area of land where precipitation falls and moves to a common outlet. As a data source for drainage, we used the vector map from the Municipal Basic Sanitation Plan [32]. This file was imported into QGIS v3.34.11 [33], where a euclidean distance map was compiled. The georeferenced leptospirosis cases were overlaid. Following this procedure, the distance values of these cases relative to the drainage network were extracted.

Rainfall data were obtained from the Hydrological Database of the Department of Water and Energy of the State of São Paulo (DAEE), through 10 meteorological stations. These stations were chosen because they had continuous daily rainfall data. After inspecting the values from the rain gauges, the flood events in the city of Campinas - SP, from 2007 to 2010, were validated through the flood database of the Civil Defense of the Municipality of Campinas. The concept of flooding used here is defined as the overflow of bodies of water (rivers, streams, lakes) due to heavy rains or an increase in water volume, which exceeds the natural limits of the riverbeds, flooding areas that are normally dry. Floods can cause significant damage to residences, infrastructure, and public safety. In urban areas, they occur where the drainage infrastructure is unable to handle large volumes of water, often due to soil impermeabilization [34]. This makes information on rain, drainage and permeability of surfaces relevant to understand disease risk for leptospirosis. Using the dates of occurrence of this type of natural disaster, the precipitation values were organized in a database.

This information was associated with the geographic coordinates of the rain gauges and, through the use of the minimum curvature interpolation method, four maps of accumulated precipitation were compiled, referring to the days on which flood events were recorded in the Campinas region during the study period (Figure S1). This method was used because it has a superior performance when compared to other interpolation methods [35].

A specific social-demographic index titled São Paulo Social Vulnerability Index (IPVS), which is based on the spatial identification of the areas of vulnerability of the resident population, classifies the urban census sectors into six categories, which range from very low or no vulnerability to very high vulnerability [36]. This index considers a range of factors influencing vulnerability, including: income, employment status, age distribution, gender, household structure, access to education, healthcare, housing, and the existence of community development projects.

Finally, the impermeable area index was calculated using the normalized difference built-up index (NDBI) derived from a Landsat TM5 satellite image from 2016 (reference ID: LT52190762010108CUB01) with a 30-meter spatial resolution. This image was acquired through a search on the image service offered by the United States Geological Survey [37]. This index was developed from information found in [38]. The NDBI is expressed by the following equation:

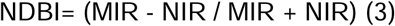

Where: MIR = mid-infrared (Band 5)

NIR =near-infrared (Band 4)

We then performed an analysis associating cases of leptospirosis with the census tracts of the municipality to verify the density of cases in each IPVS category, from extremely low to very high levels of vulnerability.

NDBI quantifies the impermeability of surfaces in a given area, considering materials such as asphalt, concrete, roofs, and other artificial covers that prevent water infiltration into the soil. This index exhibits a certain level of homogeneity, particularly in the urban center of Campinas, where its variation is limited. Therefore, for NDBI, we used the raw number of cases to examine the relationship between impermeability levels and post-flood leptospirosis risk as using case density based on NDBI could distort the identification of critical areas, as this index is not area-corrected. In contrast, for IPVS, we used case density as the response variable to indicate post-flood leptospirosis risk. Correlating case density with IPVS allows for a proportional identification of higher-risk areas, as vulnerability classes represent regions of different sizes and socioeconomic characteristics that influence exposure and interactions with the environment. All maps were developed in QGIS v3.34.11 [33].

## Results

An overview of confirmed leptospirosis cases in residents of Campinas between 2001 to 2023 revealed outbreaks consistent throughout the last decade. There were consistent outbreaks, with peaks in 2009, 2011, 2012, and 2013 (all exceeding 40 cases per year, Fig S2). After 2016, the total number of cases declined consistently, never surpassing 11 cases per year. The total number of resident cases summed 502 cases, of which 105 occurred during our investigated period (2007 to 2010). From these, 51 did not report contact with floodwater or mud (‘other risk’ category), while 54 did.

These 83 autochthonous cases (Fig. 1) were further stratified by the environment and risk exposure conditions (22 cases that reported other exposures had incomplete information). This Figure 1 illustrates the localization of the probable infection sites of leptospirosis cases.

**Fig 1.**
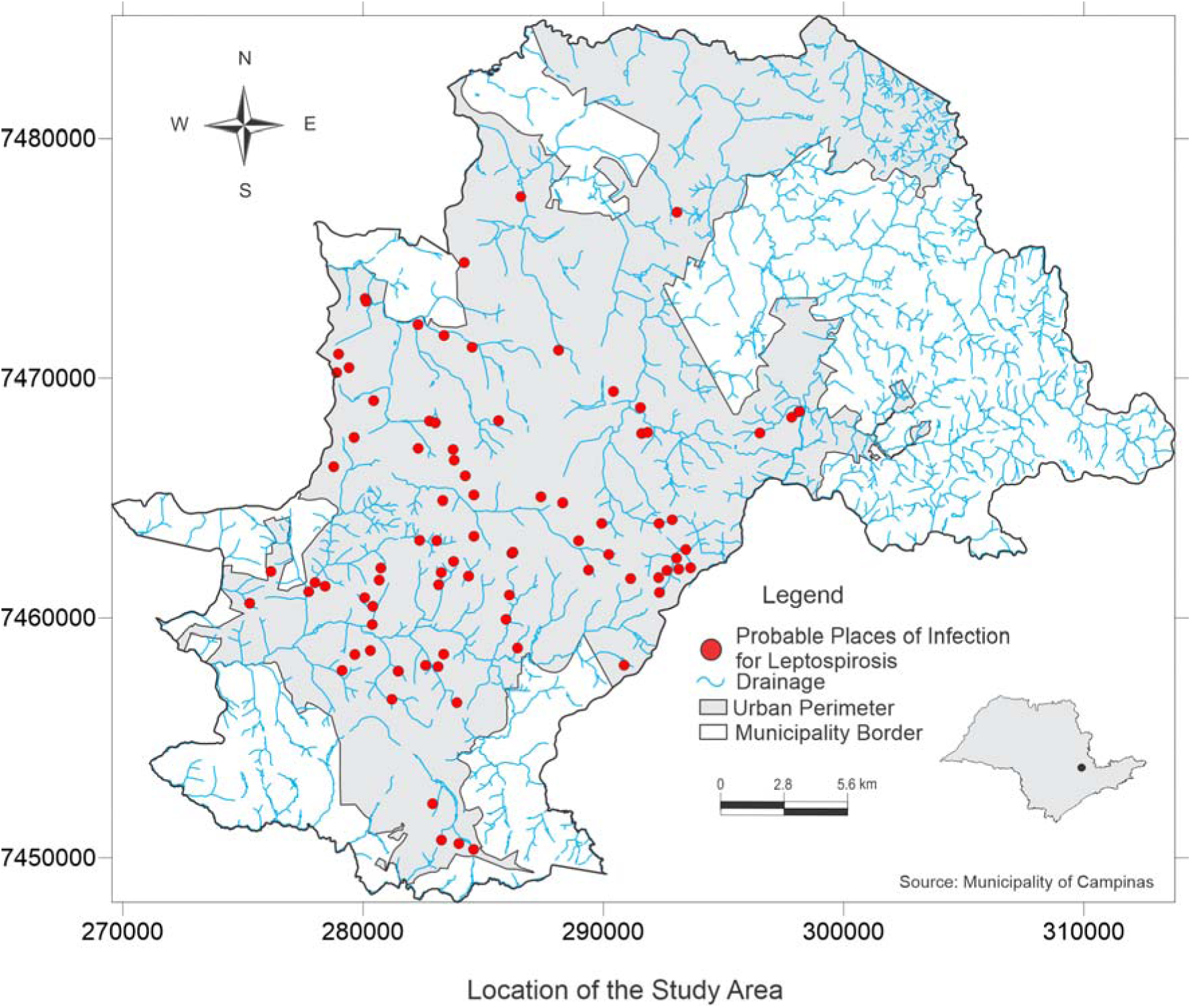
Spatial distribution of autochthonous leptospirosis cases in humans from 2007 to 2010 identified in Campinas, southeastern Brazil. Vector data (shapefile) from the Prefeitura Municipal de Campinas, as part of the city’s open data initiative. Administrative limits: https://informacao-didc.campinas.sp.gov.br/exporta_shp.php?id=119 (accessed March 2025). Streams: https://informacao-didc.campinas.sp.gov.br/exporta_shp.php?id=37 (accessed March 2025). The data is free of licences complying with the Transparência Pública Brasil (https://www.gov.br/cgu/pt-br/centrais-de-conteudo/campanhas/integridade-publica/transparencia-publica). Note that the data comes from Brazilian Public websites that may limit access to IP addresses outside of Brazil.

Fig. 2A presents the main exposures associated with the leptospirosis cases during the study period. The most significant factor was the risk related to floodwater or mud, which accounted for 51% (n=54) of the cases, while other risks represented 49% (n=51) of the cases. Contact with rodents was the second major risk factor, with 18,1 % (n = 19) cases, reinforcing the connection between leptospirosis and the presence of rats, which are the main vectors of the *leptospira* bacteria. Other risk factors include vacant lots with 8,6% (n = 9) and contact with garbage representing 6,7% (n = 7), suggesting that inadequate urban conditions may increase the risk of infection. Occupational exposure factors, such as animal farming and cleaning water tanks, account for 3.8% (n=4) and 2.9% (n=3), respectively. Factors such as grain storage, contact with crops, exposure to sewage, and bodies of water each accounted for only 1.9% of the cases (n=2 for each category), indicating that while they are potential sources of contamination, they are less frequent in the analyzed population. The category labeled "Others" had only one recorded case.

**Fig 2.**
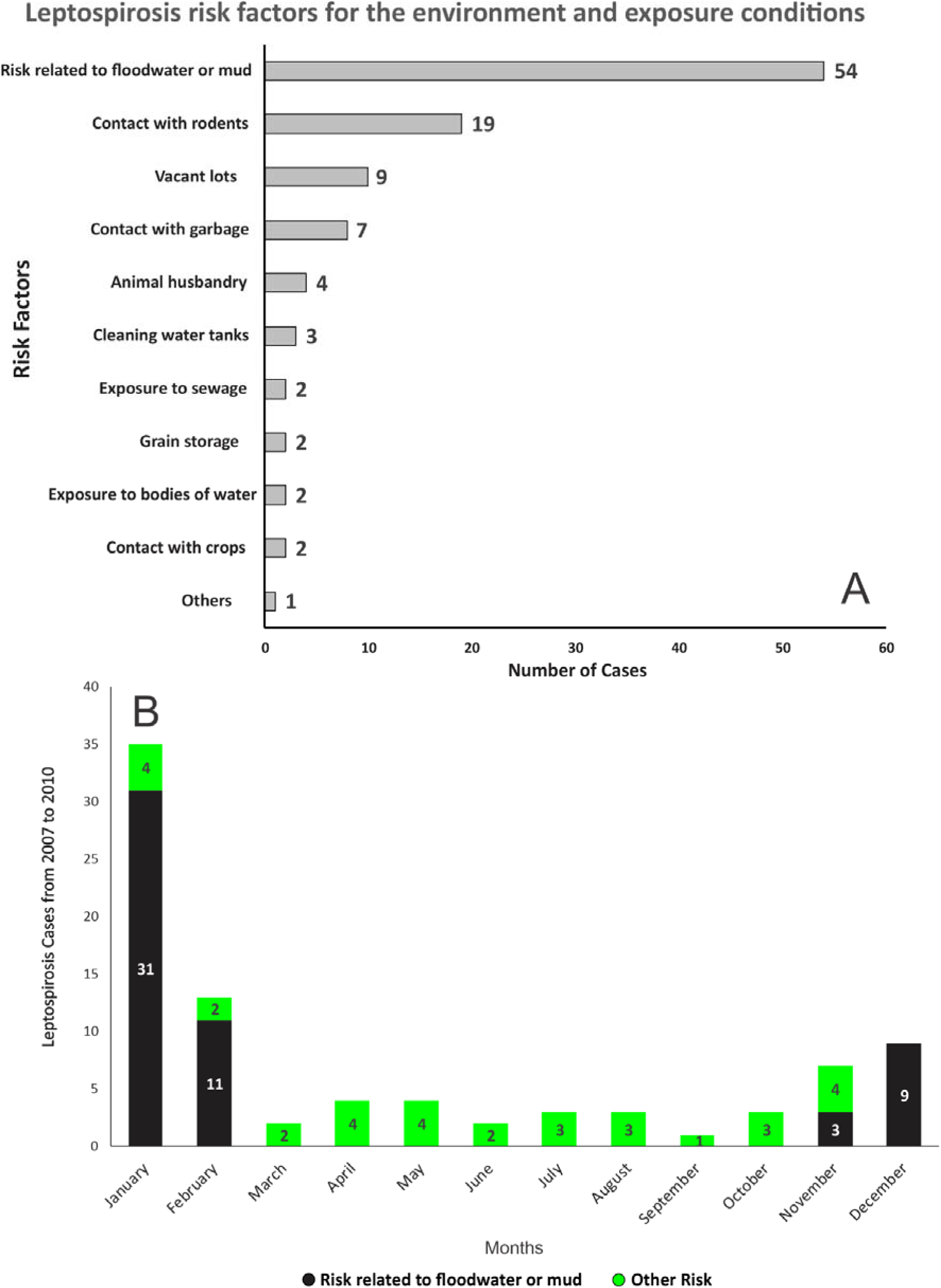
A. Exposures reported by leptospirosis cases and **B.** monthly distribution of leptospirosis cases (2007-2010) in Campinas, southeastern Brazil.

Fig. 2B shows the monthly distribution of leptospirosis cases, with the two selected categories: Risk related to floodwater or mud and Other Risks. January accounted for the highest number of leptospirosis cases, with 31 cases related to floodwater or mud. This is likely linked to heavy rains and floods that were typical for this period. In February, the number of cases related to floods decreased but remains the second highest month with 11 cases. December also shows an increase in flood-related cases, with 9 cases. In other months, the number of cases linked to floods and other risks was low, with only a few cases recorded from March to November.

The sensitivity analysis revealed a weak relationship observed between the number of cases and the amount of rainfall, suggesting that flooding, rather than rainfall alone, ncreases the risk of infection. However, positive associations between IPVS and NDBI were observed (Fig. 3A). The closer a location was to drainage systems, the higher the number of eptospirosis cases. This pattern holds true regardless of the specific mode of exposure (e.g., water/mud contact vs. other risks), as the adjusted curves for both exposure types largely overlap (Fig. 3B). Additionally, a linear relationship was observed between the frequency of eptospirosis cases and their proximity to urban drainage systems (Fig. 3B). As the distance ncreases, the frequency of infections strongly decreases (R² = 0.82), suggesting that eptospirosis cases are more concentrated near urban drainage systems. The frequency of nfections related to other risks also shows a decreasing trend but with a slightly weaker orrelation (R² = 0.79) compared to flood risk. For flood-related cases, the fitted curve is teeper, suggesting a greater impact of these factors on the risk of infection. At distances greater than 700 meters, the difference between the two types of risk decreases, suggesting hat, far from urban drains, other factors may influence the infection.

**Fig 3.**
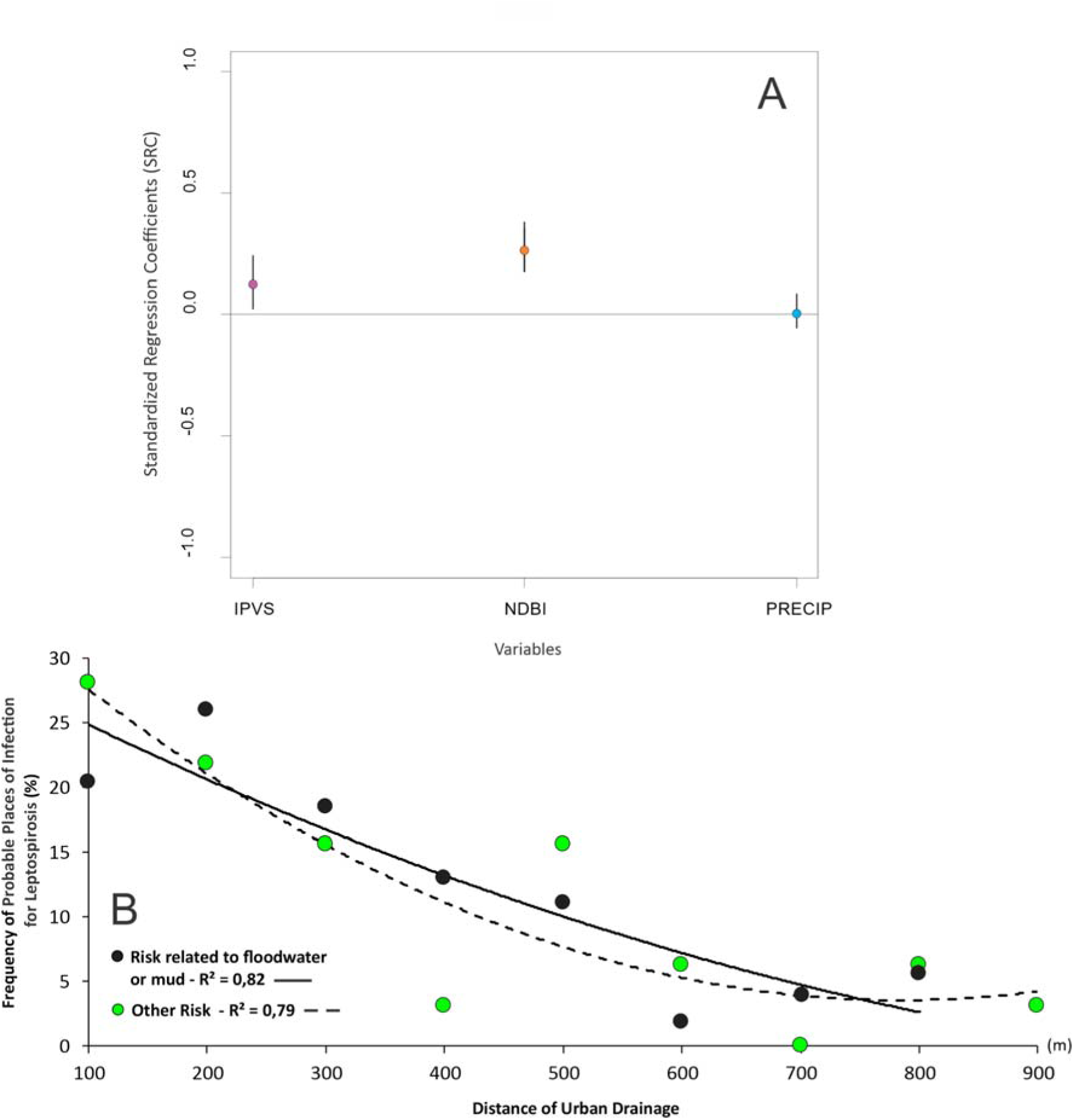
A. Sensitivity analysis of the São Paulo Social Vulnerability Index (IPVS), the Normalized Difference Built-up Index (NDBI), and precipitation, in relation to the number of leptospirosis cases associated. **B.** Relationship between the frequency of leptospirosis cases and distance from urban drainage.

The spatial distribution of leptospirosis cases overlaid with IPVS illustrates that cases are concentrated in areas of higher social vulnerability (Fig 4A), with a positive relationship between the density of leptospirosis cases and increasing levels of social vulnerability (Fig 5B).

**Fig 4.**
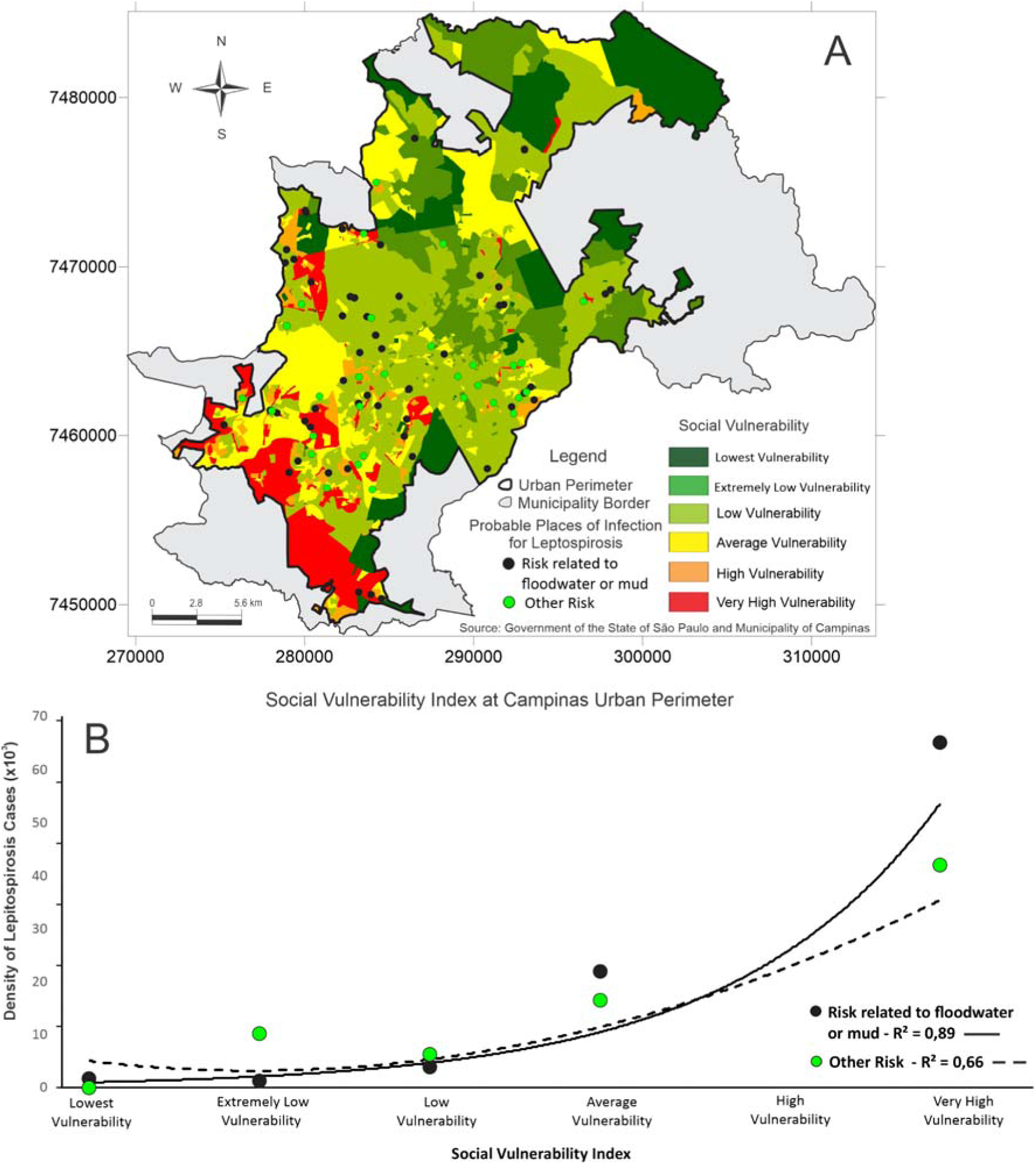
Higher social vulnerability is strongly linked to increased leptospirosis risk in Campinas, SP, Brazil. **A.** Spatial distribution of all human leptospirosis cases who reported contact with floodwater/mud overlaid with the social vulnerability (IPVS Index) from 2007 to 2010 period in Campinas, São Paulo, southeastern Brazil. B. Density of leptospirosis cases who reported contact with floodwater/mud from 2007 to 2010 according to social vulnerability levels. R² represents the proportion of variance in the response variable explained by the covariate, where values closer to 1 indicate a better fit. Vectorial data (shapefile) source from the São Paulo State Government SEADE, available at: http://ipvs.seade.gov.br/view/index.php (accessed March 2025). The data is free of licences complying with the Transparência Pública Brasil (https://www.gov.br/cgu/pt-br/centrais-de-conteudo/campanhas/integridade-publica/transparencia-publica). Note that this data comes from Brazilian Public websites that may limit access to IP addresses outside of Brazil.

**Fig 5.**
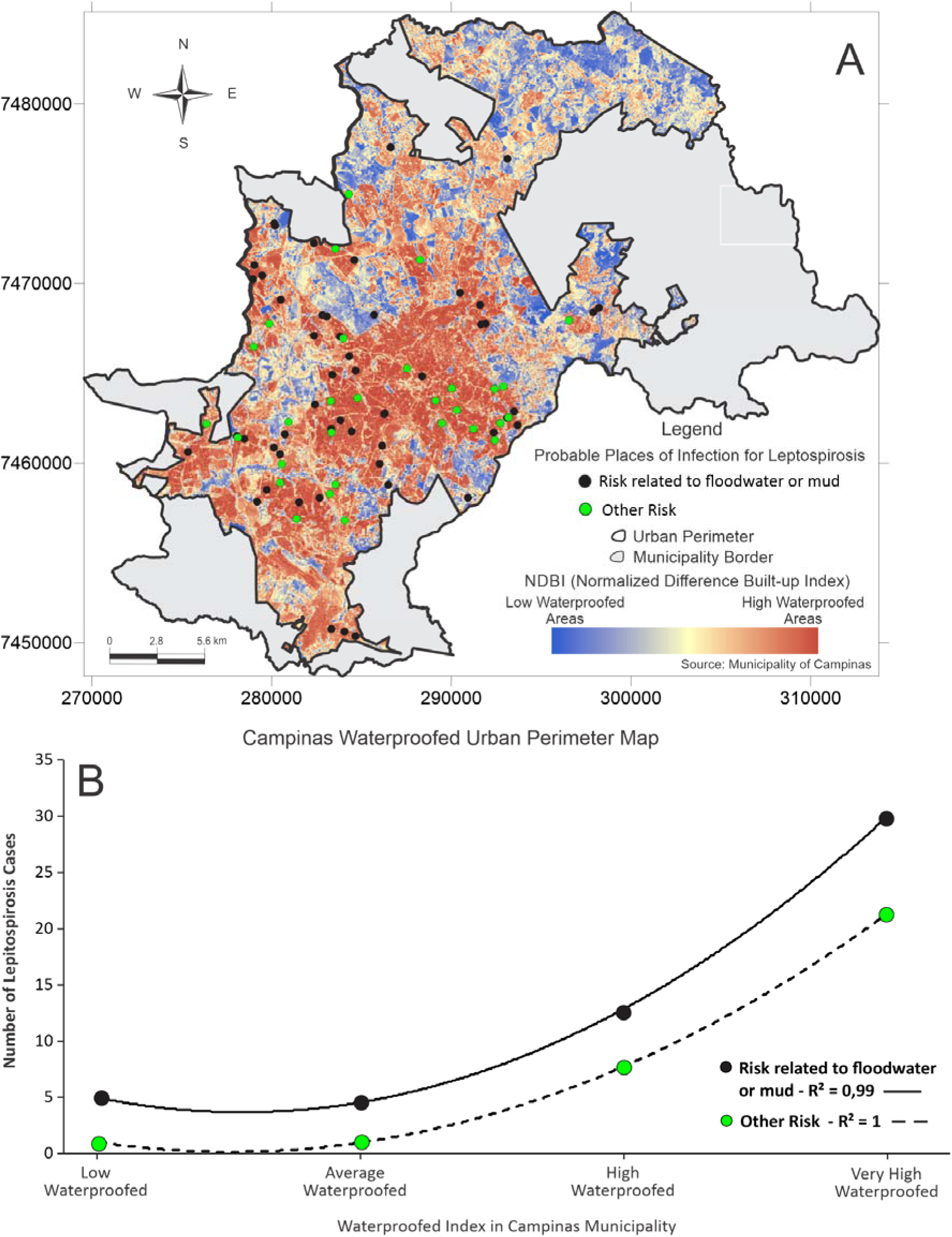
Leptospirosis cases are consistently higher in areas with more impermeable surfaces. **A.** Spatial distribution of all human leptospirosis cases who reported contact with floodwater/mud from 2007 to 2010 overlaid with ground permeability in Campinas, São

In the most vulnerable areas, cases linked to floodwater/mud exposure outnumbered cases from other exposures, suggesting that flooding-related contamination disproportionately affects the most vulnerable populations (Fig. 4). As the IPVS increases (indicating higher social vulnerability), the density of leptospirosis cases related to floodwater/mud exposure shows a marked increase (Fig. 4B). The relationship is non-linear and exhibits a steep rise, especially in regions with higher vulnerability, implying that communities with greater social vulnerability are more likely to be exposed to flood-related leptospirosis. The curve suggests that as vulnerability increases, flood-related risks and thus leptospirosis cases significantly increase as well, with a strong correlation (R² = 0.88). The density of leptospirosis cases attributed to other risks (such as rodent contact or poor urban conditions) also increases with higher vulnerability, but the trend is more gradual, indicating that other social determinants, like sanitation or living conditions, contribute to the risk of leptospirosis, but at a slower rate compared to the flood-related risk. This correlation is slightly weaker than that of flood-related cases (R² = 0.64), suggesting that although vulnerability impacts both types of risks, the connection is stronger for cases related to floods.

The spatial distribution of human leptospirosis cases in relation to ground permeability in Campinas show that cases predominantly occurred in areas with highly impermeable grounds (Fig 5A). This emphasizes the role of urban environmental characteristics in disease transmission, likely due to the accumulation of floodwater and subsequent exposure risk (Fig 5B). However, unlike the drainage-related cases, the two exposure curves (floodwater/mud *versus* other risks) do not overlap but follow the same trend, with floodwater/mud exposure consistently leading to more cases. As the Waterproofed Index increases (indicating better waterproofing), the number of leptospirosis cases linked to floodwater/mud exposure increases significantly. The relationship is non-linear and shows a sharp upward trend as the waterproofing level becomes higher. This suggests that areas with better waterproofing are correlated with more flood-related leptospirosis cases, which may imply that, even in better-drained areas, there may be floods to expose individuals to the risk of leptospirosis. This correlation is strong (R² = 0.87), indicating that the higher the levels of waterproofing, the more concentrated the leptospirosis cases linked to floods are in these areas. The other risk category also increases with higher waterproofing, but the trend is less steep. This suggests that while there are more leptospirosis cases attributed to other factors (such as rodent exposure, unsanitary conditions, etc.) in areas with higher waterproofing, the change is not as pronounced as in flood-related cases.

Paulo, southeastern Brazil. **B.** Number of leptospirosis cases who reported contact with floodwater/mud from 2007 to 2010 according to the waterproof index. Vector data (shapefile) from the Prefeitura Municipal de Campinas, part of the city’s open data initiative.

Administrative limits: https://informacao-didc.campinas.sp.gov.br/exporta_shp.php?id=119 (accessed March 2025). The data is free of licences complying with the Transparência Pública Brasil (https://www.gov.br/cgu/pt-br/centrais-de-conteudo/campanhas/integridade-publica/transparencia-publica). Note that this data comes from Brazilian Public websites that may limit access to IP addresses outside of Brazil.

## Discussion

Here, we discuss insights from our findings around leptospirosis cases and temporal trends in Campinas, and discuss its relevance for global combat to leptospirosis through urban planning and social development. There is a clear connection between leptospirosis cases and social vulnerability which underpin inequalities in access to services, infrastructure quality and exposure to environmental hazards. Leptospirosis risk was amplified post-flooding, particularly for cases residing in areas of high social vulnerability and close to highly impermeable areas, when compared to cases who did not report exposure to floodwaters or mud. In our analysis, we quantified these connections, highlighting a disproportionate increase in leptospirosis risk for vulnerable conditions.

The monthly variation of cases show that flood-associated leptospirosis cases are more prevalent during the rainy season and flood periods, with peaks in January and February, and a slight increase in cases in December (Fig. 2B). On the other hand, cases associated with other risks seem to be less seasonal and more evenly distributed throughout the year, indicating that these may have a more dispersed distribution in the urban environment. While this pattern is expected given leptospirosis’ waterborne nature, our analysis provides a formal quantification of these dynamics, reinforcing the clear trends with empirical data. This not only validates common knowledge but also offers a measure of the interplay between flooding and disease risk.

We identified a clear decrease in the probable location of leptospirosis infection as the distance from drainage increases, with the peak in cases observed at 200 m (Fig. 3B). This is likely due to floodwaters accumulating closer to overflowing drains, increasing the risk of exposure to the *Leptospira* bacteria. This peak makes sense from the point of view of rodent host habitat and movement behaviour. Rodents are associated with roosts around dirt banks close to drainage and sewer, likely 3x more common to be found in areas of bare soil or mixed ground [39], and those areas are likely to be right next to paved/closed drainage (proximity to drainage = 0) in our context.

As impermeability increases, leptospirosis cases also rise, indicating that impermeable surfaces may favor the accumulation of contaminated rainwater, contributing to outbreaks of the disease (Fig. 5B). Therefore, preventive measures such as improvements in sanitation, rodent control, and proper waste management are essential to reducing the incidence of the disease. Rapid urbanization and unplanned growth has resulted in the occupation of environmentally sensitive areas, increasing the exposure of vulnerable communities to floods and the risk of leptospirosis [40]. In Campinas, these situations, combined with the presence of rivers such as the Atibaia and Capivari, have become a recurring event, especially during the rainy summer season (December to March). The epidemiological data showed a significant increase in leptospirosis cases following periods of heavy rainfall, where low-income regions are more susceptible to flooding, and consequently, leptospirosis outbreaks [41]. Particularly in extensive areas of Campinas, systematic processes of urban planning that lead to people being exposed to the hazard are inferred, with persistent floods, floodplains management and stormwater design being below optimal conditions [42], with numerous plans on paper, yet lacking alignment with real-world needs [43]. Plans for managing can also be funded by different government levels, but prefecture and state level funding of different infrastructure are not necessarily being conducted in an integrative way. Specifically for the eastern regions of Campinas (Cabras), it is not clear which standards are considered flood prevention for pedestrians. Therefore, the post-flood transmission risk of leptospirosis in Campinas remains a health problem since the last decades [41]. These risk areas for flood-prone leptospirosis cases are closely related to high-social vulnerability (Fig. 4B). More than half a million people (∼60% of the total population of Campinas) live in areas of high and medium-high vulnerability and most of the population in these areas have an average income of between one and two minimum wages. These areas are more clustered in the floodplains of the Quilombo and Atibaia Rivers [44] .

Environmental characteristics of urban areas such as populated urban slum areas that lack adequate water supply and sanitation [45] can significantly contribute to disease risk. Other key elements include habits and hygiene behavior, as well as ecosystem and pathogen-related factors. When these factors come together, they create a permeable proximity with the pathogen in the environment [14]. The difficulty in assessing the influence of environmental conditions on the rapid dissemination of infection is that these conditions vary at different time intervals and at different scales. The severity and frequency of rainfall, the compaction degree of the urban areas, and the quality of sanitary infrastructure are examples of circumstances which vary spatially and socially [15,46]. Assessing these conditions can be more difficult in areas where flooding events are frequent. Finding consistent patterns in areas with a higher incidence of leptospirosis often coincide with regions of high population density and low urban infrastructure index [47] is useful for preparedness plans informed by spatial indexes and remotely sensed data [48]. Neighborhoods with specific characteristics, such as proximity to streams and irregular occupation of risk areas, with the highest levels of built areas denoting higher impermeability, present a higher concentration of cases. It is exacerbated in neighborhoods such as Campo Belo and the Sousas, which often face seasonal floods that increase population exposure to contaminated water.

In Brazil, 10% of leptospirosis cases evolve to high severity. Leptospirosis remains a threat despite the recent decrease in the total number of cases, with the number of deaths breaking records in 2023 [49], and the outlook not looking positive for the 2024 flood season. For people at risk in areas that have precarious drainage infrastructure (such as Princesa d’Oeste, Amoreiras Avenue surroundings), it is essential for governance to prioritize the allocation of resources toward awareness campaigns and mitigation strategies aimed at reducing the impact of leptospirosis during flooding events.

In contexts similar to the one we investigated here, local authorities should consistently invest in education and early warning systems (such as the ones proposed for malaria combat [50,51]) that may inform stakeholders about the areas associated with flooding and leptospirosis transmission, as well as the preventive measures they can take. Furthermore, fostering robust community networks is crucial for developing effective response plans to address flooding scenarios, which are projected to increase in the Global South [52]. These plans should include strategies for supporting individuals who may need to commute or find shelter when floods threaten their homes, ensuring that vulnerable populations have access to safe spaces and necessary resources during emergencies. By strengthening community resilience and enhancing awareness, governance can significantly mitigate the health risks associated with leptospirosis in flood-prone areas.

Climatic extremes [53] are driven by multidimensional environmental factors that in turn interact with other risk components influencing risk of transmission of leptospirosis [54]. Because extreme events can continuously disturb and affect urban regions (like the proposed ‘Joseph effect’ in Campinas) [55], mitigation action plans are probably more needed than ever. São Paulo State has been suffering from higher extreme climatic events clustering in shorter rainy seasons and longer periods of heatwaves and drought [56]. Retrospective analysis like the one we present are useful, especially when trends are pervasive like we see with leptospirosis and floods in Campinas. Post-flood cases are common in urban settings, where a patient can be exposed to multiple contaminated water sources in a very brief period of time [57]. There is a current gap in studies about urban drainage network adaptation to climate change [58]. In urban settings in Brazil, infected rats with a high prevalence of infection were found among all age classes indicating efficient transmission [2,59]. Sustainable rodent management options should be explored [60]. Given the complexity of factors contributing to leptospirosis transmission during floods, preventive strategies must be multisectoral [61], involving sanitation, rodent control, and public awareness. In Campinas, Epidemiological Surveillance and Civil Defense play an essential role in the prevention and response to leptospirosis outbreaks during flood events. Measures such as strengthening drainage infrastructure, adequate solid waste management, and educational campaigns are essential to reduce population exposure to contaminated water. Actions like these have been implemented in Campinas, but effectiveness is limited by the need for broader investments in urban infrastructure and services [17].

Geographical studies on leptospirosis are essential for epidemiological surveillance, as they help identify spatial patterns, risk areas, and environmental factors influencing disease spread. GIS tools enable visualization of case distribution and correlations with factors like flooding, drainage proximity, and urbanization, supporting targeted interventions and resource allocation. These approaches also aid in monitoring outbreaks, planning preventive measures, and improving responses to health emergencies by guiding surveillance efforts and public policies. Ultimately, spatial analysis can greatly benefit disease monitoring and contribute to strategies that reduce transmission, particularly in vulnerable communities.

### Study limitations

There are several limitations to conducting neglected disease research in Brazil. The data associated with patients in the SINAN system is often incomplete, particularly regarding exposure history, such as the probable site of infection, the history of contact with floodwater or mud, or other environmental factors. Here, we noted that the databases contained typographical errors, missing addresses of the probable infection site, or, when present, they were incomplete or incorrect. For example, of the 51 cases that reported Other exposure risks, only 19 had complete information in the database. Leptospirosis may be broadly underreported in Brazil [62]. This can occur due to a lack of diagnosis, insufficient resources for data collection, or a shortage of trained healthcare professionals. Additionally, as reported here, the quality of the data collected and recorded in SINAN can vary depending on the training of healthcare professionals and the resources available and other limited enabling conditions across different regions. This can lead to errors in completing the notification forms and affect the accuracy of the information. Another limitation is the demand for other indicators, such as rodent density or an infestation index. Such information could improve the effectiveness of the associations between leptospirosis and exposure environments. Moreover, it is important to highlight that some environments not explicitly categorized as “mud contact” in the SINAN forms, such as vacant lots, crop fields, and animal husbandry sites may potentially involve exposure to mud. These areas are typically unpaved and can accumulate water and organic waste, especially during or after rainfall, irrigation, or animal activity, resulting in muddy soil conditions favorable to Leptospira survival. The absence of direct attribution to mud may underestimate the general exposure pathways. This limitation reinforces the need for complementary field investigations and broader environmental characterization when analyzing leptospirosis risk factors.

## Conclusion

This study underscores the spatial patterns of leptospirosis in Campinas, with a particular focus on how socio-environmental and climatic factors influence the disease’s risk. Proximity to drainage systems, impermeable surfaces, and high social vulnerability emerge as key determinants of case distribution, with flood-prone areas experiencing the greatest risk, which corroborates our hypothesis. Cases associated with floodwater and mud exposure are consistently more frequent in the most vulnerable communities, underscoring the inherently compounded risk posed by inadequate infrastructure and environmental conditions. The increase in extreme weather events worsens the risk of contamination by increasing the frequency of floods, emphasizing the need for effective preventive measures and public policies. The study suggests that improvements in urban infrastructure, including drainage systems, access to basic sanitation and flood response systems, could significantly reduce cases of leptospirosis particularly in the Global South. Furthermore, collaboration between different government sectors, such as health, environment, and urban planning, remains essential to mitigate the impacts of floods and control the transmission of the disease. The study also highlights the importance of educational campaigns to raise awareness about the risks of leptospirosis and protective measures during flood periods. The methods used can be applied to other cities with similar socioeconomic and environmental contexts, aiding in the generation of evidence that could inform the development of public health policies aimed at reducing leptospirosis.

## Supporting information

**S1 Fig. Spatial distribution of rainfall events from 2007 to 2010 that caused flooding events in Campinas, São Paulo, southeastern Brazil.** Administrative limits: https://informacao-didc.campinas.sp.gov.br/exporta_shp.php?id=119 (accessed March 2025). The data is free of licences complying with the Transparência Pública Brasil (https://www.gov.br/cgu/pt-br/centrais-de-conteudo/campanhas/integridade-publica/transparencia-publica). Note that this data comes from Brazilian Public websites that may limit access to IP addresses outside of Brazil.

**S2 Fig. Confirmed leptospirosis cases in Campinas, southeastern Brazil between 2001 and 2023.**

## Acknowledgments

TSA thanks the Epidemiological Surveillance of Campinas Municipality for providing the leptospirosis data. RLM is supported by an Australian Research Council Australian Laureate Fellowship (FL240100037) funded by the Australian Government. RLM thanks the Massey University Foundation and the Morris Trust for their support. The funders had no role in study design, data collection and analysis, decision to publish, or preparation of the manuscript.

