## Supporting Information for "Leptospirosis in Campinas, Brazil: The interplay between drainage, impermeable areas, and social vulnerability"

#### **This file includes:**

Supplementary Figures 1-2

**S1 Figure.** Spatial distribution of rainfall events from 2007 to 2010 that caused flooding events in Campinas, São Paulo, southeastern Brazil. Administrative limits: <https://informacao-didc.campinas.sp.gov.br/exporta_shp.php?id=119> (accessed March 2025). The data is free of licences complying with the Transparência Pública Brasil (<https://www.gov.br/cgu/pt-br/centrais-de-conteudo/campanhas/integridade-publica/transparencia-publica>). Note that this data comes from Brazilian Public websites that may limit access to IP addresses outside of Brazil.

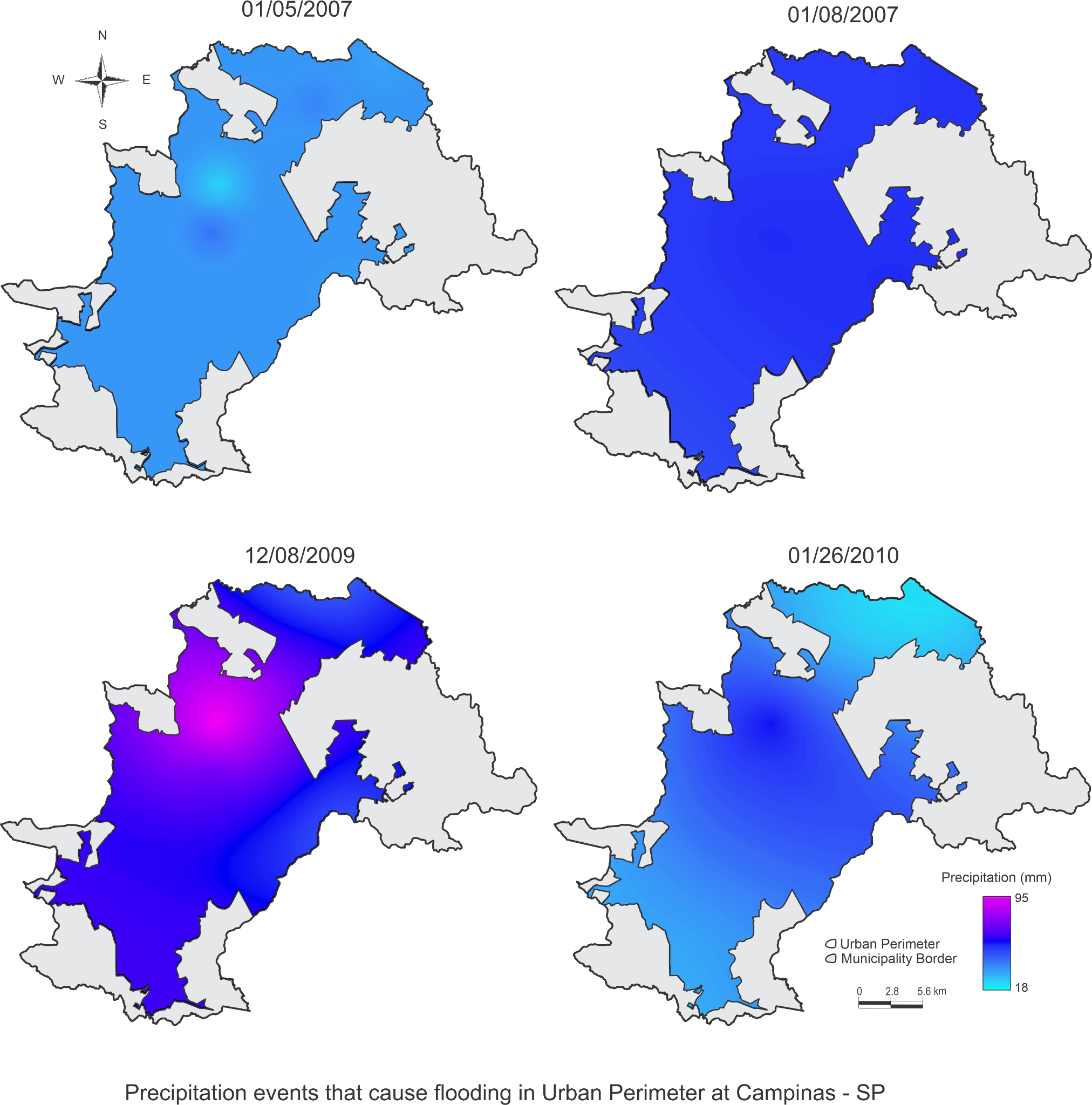

**S2 Figure.** Confirmed leptospirosis cases in Campinas, southeastern Brazil.
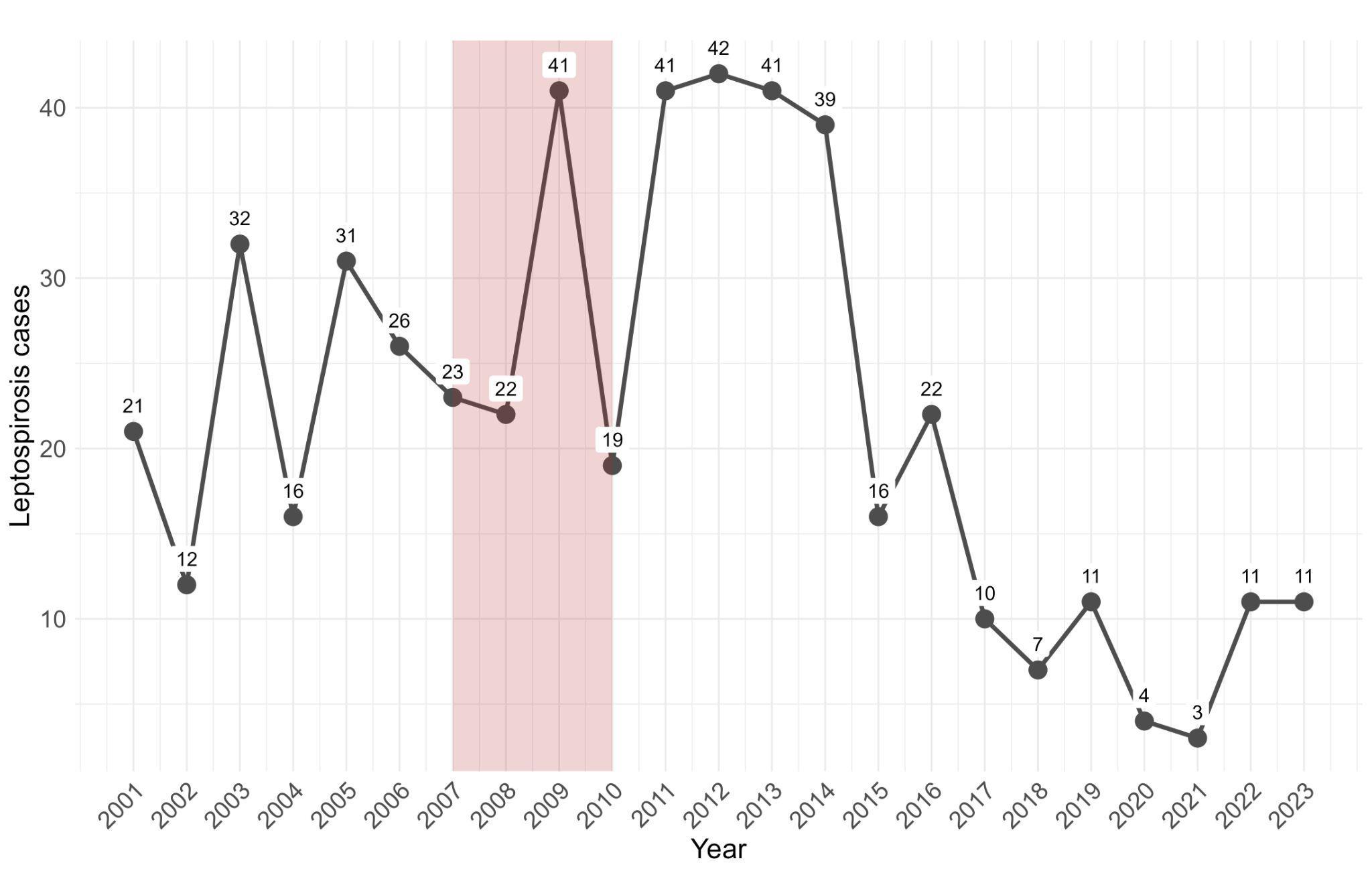

Source: Datasus (2024).
